# SOD1 Enzymatic Activity in CSF from ALS patients with and without *SOD1* mutations

**DOI:** 10.1101/2024.06.15.599161

**Authors:** Laura Leykam, Karin M.E. Forsberg, Ulrika Nordström, Karin Hjertkvist, Agneta Öberg, Eva Jonsson, Peter M. Andersen, Stefan L. Marklund, Per Zetterström

## Abstract

Superoxide dismutase-1 (SOD1) mutations are a common cause of amyotrophic lateral sclerosis (ALS). Intrathecal gene therapy using the antisense-oligo-nucleotide drug tofersen to reduce *SOD1* expression shows significant effects on disease progression and has recently been approved in the United States and the European Union. However, the discovery of children homozygous for four different inactivating *SOD1* mutations developing the Infantile SOD1 Deficiency Syndrome (ISODDES) with injury to both upper and lower motor systems suggests that low SOD1 activities may be deleterious in humans. Monitoring SOD1 activity in cerebrospinal fluid (CSF) from tofersen-treated patients is advisable but difficult due to low levels and the presence of the isoenzyme SOD3. We here present a method to efficiently remove SOD3 from CSF using highly specific immobilized antibodies and subsequent measurement of the SOD activity as a sensitive assay for the SOD1 activity in CSF. We applied the method on CSF samples from ALS patients and controls and used paired erythrocyte samples for comparison. In ALS patients with wildtype *SOD1,* the SOD1 activity in CSF was the same as in controls, but patients with mutant *SOD1* show lower activity in CSF, even patients with mutants previously reported to have full activity in erythrocytes. Activity variation is large among patients carrying the same *SOD1* mutation and larger than seen in erythrocytes and in post-mortem central nervous system tissue. SOD1 in CSF shows a high specific activity, indicating that it is mainly native. Lastly, we identified a discrepancy between the SOD1 activity and protein level measured with ELISA in both CSF and erythrocytes. Since antibodies for SOD1 ELISA-quantification are raised against the native wildtype enzyme, the content of mutant SOD1s may be underestimated. Direct analysis of SOD1 enzymatic activity in CSF is therefore a more reliable way to monitor the effect of SOD1-lowering compounds.

## Introduction

Amyotrophic lateral sclerosis (ALS) is a heterogeneous neurodegenerative disorder characterized by adult-onset progressive degeneration of motor neurons resulting in atrophy of skeletal muscles and inevitable death, usually within 1-5 years from onset (Brown & Al-Chalabi, 2017; Masrori & Van Damme, 2020). ALS is classified into sporadic (idiopathic) disease with unknown etiology constituting the majority of patients but a third of all patients carry pathogenic mutations in some 40 different genes, the most common in Europe being *C9orf72, SOD1* and *FUS* (Brown & Al-Chalabi, 2017). Mutations in the gene encoding superoxide dismutase-1 (*SOD1,* EC 1.15.1.1) is globally the most common cause of ALS known, accounting for 1-5% in different populations (Rosen, 1993). SOD1 is expressed in all cells in the human body and plays an essential role as a radical scavenger by neutralizing the superoxide anion radical to hydrogen peroxide and oxygen (Eleutherio et al., 2021).

SOD1 is located mainly in the cytosol but also in the nucleus and mitochondrial intermembrane space and constitutes around 0.1% of the protein in the human brain (Forsberg et al., 2010). Two other SOD isoforms, SOD2 found in the mitochondrial matrix (Weisiger & Fridovich, 1973) and the extracellular superoxide dismutase (SOD3, EC 1.15.1.1) (Marklund, 1982), are also expressed in humans. While no mutations in SOD2 or SOD3 have been associated with ALS, since 1993 some 234 coding mutations in the small *SOD1* have been associated with causing ALS usually as a Mendelian dominant trait, but recessive inheritance and *de novo* mutations have also been found (Andersen, 2006; Müller et al., 2022). In erythrocytes, most mutations result in a reduction in the enzymatic activity to various degrees with many mutants that lack activity having, on average, near half the normal SOD1 activity found in control individuals (Saccon et al., 2013). Two mutants (G127X and G93R) however have been proposed to have a dominant-negative effect (Andersen et al., 1997; Orrell et al., 1995). Conversely, seven mutations (C6S, E40G, D90A, D109Y, E100K, L117V, and G133A) have been reported to show essentially normal enzymatic activity in erythrocytes from patients (Keskin et al., 2017). This, together with the finding that transgenic mice overexpressing human SOD1-mutants develop a murine ALS-like disease (but have preserved normal function of their own murine SOD1) and knock-out of *SOD1* in mice does not cause an ALS-like phenotype, have fueled the prevailing toxic gain-of-function theory (Andersen et al., 1995; Gurney et al., 1994; Jonsson et al., 2006; Reaume et al., 1996). Also, expression of a human dismutase-inactive variant (G85R) in mice results in the same ALS phenotype and survival independently of the mice expressing the endogenous mouse SOD1 gene or not (Bruijn et al., 1998) further uncoupling SOD1 enzymatic activity from ALS pathogenicity. The collective data proposes a hypothesis involving destabilization of the SOD1 monomer, misfolding and increased aggregation of SOD1 in the cytosol of motor neurons in a dose-dependent manner (Kato et al., 2000). Spinal cord inoculation experiments in mice suggest that the human SOD1 aggregates are ALS-transmitting prions (Ayers et al., 2014; Bidhendi et al., 2016; Ekhtiari Bidhendi et al., 2018) and so far, two different tertiary structures associated with different phenotypes in mice models have been reported (Bergh et al., 2015). The conformation and dose-dependent formation and propagation of human SOD1 prions could be the core pathological mechanism in SOD1-linked ALS.

Use of either antisense oligonucleotides (ASO) or siRNA compounds to reduce the expression of SOD1 offers new potential treatments for ALS. A number of drug trials have been performed, first in rodent models and later in symptomatic ALS patients with *SOD1* mutations (Borel et al., 2016; Miller et al., 2020; Miller et al., 2022; Miller et al., 2013). In April 2023, the first ASO for ALS (tofersen (Qalsody)) was approved by the U.S. Food and Drug Administration for the treatment of ALS patients with mutations in *SOD1* (Blair, 2023) and in May 2024 the European Commission did the same. Designed to initiate SOD1 mRNA degradation, tofersen is non-mutant allele specific and reduces the expression of both the mutant and the non-mutant SOD1 protein in the order of 29-40% when given at a dose of 100 mg intrathecally every four weeks (Miller et al., 2022). Since 2022, a compassionate/Early Access Program of tofersen is ongoing in some countries and the treated patients are being closely monitored. The first reports confirm that tofersen is safe and treatment is in many patients associated with a significant slowing of disease progression as measured by the ALS Functioning Rating Scale Revised, slowing the decline of respiratory function, and reduction in neurofilament L and H biomarkers (Forsberg et al., 2024; Meyer et al., 2023; Wiesenfarth et al., 2024). Currently, tofersen is also evaluated for presymptomatic efficacy in carriers of pathogenic *SOD1* mutations in the ongoing ATLAS trial (NCT04856982) (Benatar et al., 2022). In none of the published studies have the SOD1 enzymatic activity or mutant SOD1 protein levels been assessed.

During the conception of SOD1-protein knock-down studies in humans, the question of possible negative effect(s) of intermittent and/or chronic SOD1 protein reduction was raised (Miller et al., 2013), in particular since the intervention will be lifelong. Hence, foremost for safety reasons, it is important to measure SOD1 enzymatic activity in the cerebrospinal fluid (CSF) when evaluating SOD1-expression reducing treatments. Also, a threshold below which activity loss leads to negative effects should be defined. Performing such measurements in CSF is challenging since the SOD1 activity in CSF in healthy subjects is low (Jacobsson et al., 2001; Marklund et al., 1982) and due to the presence of the SOD3 isoform coincident with SOD1 (Marklund et al., 1982).

Here we present a novel method to specifically measure SOD1 activity in CSF samples from ALS patients and controls without interference of SOD3. We also correlate these values with the SOD1 protein content and show that *SOD1* mutations have a different effect on SOD1 activity in CSF compared to erythrocytes. We suggest that SOD1 activity should be analyzed in CSF to evaluate the effect of *SOD1* knock-down.

## Materials and Methods

### Patients and control participants

Participating patients were seen at the Department of Neurology, Umeå University Hospital. The ALS patients were diagnosed according to the European Federation of Neurological Societies diagnostic algorithm for managing ALS (Andersen et al., 2012). CSF was collected as part of the diagnostic work up procedure or, rarely, if the patient participated in other studies or came for second-opinion evaluation. Typically, the spinal tap was done non-fasting and in the morning. The level of the spinal tap was usually L3-L4 or L4-L5 and an atraumatic needle was used. The first collected CSF was used for the diagnostic tests, but an additional 4-8 mL of CSF was collected and stored for possible later diagnostic analysis. If additional clinical analysis was not warranted, the CSF could, with consent from the patient and the ethical committee (FEK 1994 dnr 94-135 with later amendments adhering to the Declaration of Helsinki (WMA 1964), used for medical research purposes. The standard procedure is to collect the CSF by drip into a polypropylene tube, aliquot the CSF into 1 or 2 ml tubes that are then immediately frozen to –80°C without prior centrifugation. For blood enzymatic and DNA analysis, with a separate written informed consent, venous blood was drawn into ethylenediaminetetraacetic acid (EDTA)-containing tubes and immediately centrifuged and separated into plasma, erythrocytes, and buffy coat as described (Keskin et al., 2017).

Mutation analysis of *SOD1* and a number of other ALS-causing genes were performed as described (Müller et al., 2022). Samples were collected between 1995 and 2023. The cohort consisted of 45 ALS patients with pathogenic mutations in *SOD1*, 19 patients with ALS with a pathogenic hexanucleotide repeat expansion in *C9orf72*, 26 apparently sporadic ALS patients (all with a negative family history and screened for ALS genes), as well as 85 control subjects. The control subjects were patients with a variety of non-motor neuron disease diagnosis seen at our department, most common diagnosis were headache, paresthesia and polyneuropathy (summarized in Table S1). The spinal tapping, collecting and storage procedure for the control samples were identical to the ALS patients. Since *SOD1* mutations have widely different effects on SOD1 enzymatic activity in blood (Andersen et al., 1995; Borchelt et al., 1994; Bowling et al., 1995; Deng et al., 1993; Jonsson et al., 2002; Keskin et al., 2017; Marklund et al., 1997), we chose to stratify the patients carrying *SOD1* mutations into stable and unstable depending on the erythrocyte SOD1 activity. The cut-off used was arbitrarily set at the lowest value detected for any patient homozygous for the D90A mutation known to have wildtype like activity in erythrocytes and tissues (Andersen et al., 1995; Jonsson et al., 2002).

### Handling of CSF samples from the biobank

CSF sample tubes were removed from the –80°C freezer storage and immediately transferred into a styrofoam box containing dry ice. Samples were thawed for 5 min in a water bath at 24°C and CSF aliquoted into low protein binding tubes. Aliquoted and original sample tubes were quickly frozen on dry ice before storing them at –80°C. The handling of the CSF samples at room temperature (RT, 22°C) was limited to a maximum time of 20 min.

### Total protein quantification

The total amount of protein present in the samples was quantified using the Bio-Rad Protein assay (Bio-Rad Laboratories, Hercules, California, USA), following the standard protocol. A 4 mg/mL bovine serum albumin stock solution in 0.9% NaCl stored at –80°C was used as a protein standard. CSF samples were diluted 1:5 with 1x phosphate buffered saline (0.137 M NaCl, 2.7 mM KCl, 4.3 mM disodium hydrogen phosphate and 1.4 mM potassium dihydrogen phosphate). 20 µL of protein standard and CSF samples were mixed with 200 µL of Ultrapure water (Mili-Q water Purelab flex2, VWS, High Wycombe, UK) and 1:5 diluted dye reagent pipetted in separate wells in a standard 96 well microtiter plate (Thermo Fisher Scientific, Waltham, USA). The plate was immediately mixed and measured at 595 nm using the Milenia Kinetic Analyzer (Diagnostic Product Cooperation, Los Angeles, California, USA).

### Antibodies

Anti-SOD3 antibodies were raised in rabbits against full length human SOD3 purified from human lung tissue as previously described (Marklund, 1984). Anti-SOD1 antibodies were raised using full-length native human SOD1 purified from erythrocytes (Marklund et al., 1997) as antigen in rabbits as previously described (Zetterstrom et al., 2011) and also in chickens with the native SOD1 coupled to keyhole limpet hemocyanin and affinity purified for IgY. Additionally, a peptide corresponding to amino acids 24-39 in human SOD1 was used as antigen to immunize rabbits as described (Jonsson et al., 2004). The antibodies were purified as previously described (Zetterstrom et al., 2011).

### Immunocapture and depletion of SOD isoenzymes from CSF

The SOD specific antibodies were coupled to CNBr-activated Sepharose Fast Flow (GE Healthcare) following the manufacturer’s protocol at a concentration of 5 mg/mL wet gel. The coupled gel beads were stored in 0.1 M Tris-HCl buffer with 0.5 M NaCl at pH 8 and 0.02% sodium azide. Before use, the gel beads were washed five times with coupling buffer (0.1 M sodium hydrogen carbonate pH 8.3 containing 0.5 M NaCl), collected by centrifugation at 500 rpm for 2 min at RT and the supernatant removed. After the last wash, buffer was left to create a 50% bead slurry. CSF samples were thawed at RT, mixed thoroughly, and placed on ice. 400 µL CSF was transferred to a fresh 2 mL reaction tube with flat bottom and 20 µL of 50% bead slurry was added. Tubes were transferred to an Intelli-Mixer (ELMI, Riga; Latvia) incubating for 2 hours at 4°C using the program U at 50 rpm. The remaining CSF sample was stored on ice during the whole procedure. After 2 hours, the beads were pelleted by centrifugation at 3000 rpm for 5 min at 4°C. 300 µL of the supernatant was carefully removed and transferred into a fresh 1.5 mL reaction tube for further analysis without disturbing the sedimented beads.

### Immunoblotting

The Western blots were carried out as previously described (Jonsson et al., 2004; Zetterstrom et al., 2011) but with Criterion TGX Stain-Free Any-KD precast gels (Bio-Rad), Trans-Blot Turbo Midi Nitrocellulose 0.2 µm transfer packs (Bio-Rad), and a Trans-Blot Turbo apparatus (Bio-Rad) using the 7 min turbo protocol for midi gels. CSF samples were diluted 1:2 with 2x sample buffer for sodium dodecyl sulfate–polyacrylamide gel electrophoresis. The 24-39 anti-peptide SOD1 antibody and the anti-SOD3 antibody were both used at a concentration of 1 µg/mL. The chemiluminescent reagent was ECL (GE Healthcare) and the ChemiDoc Imager (Bio-Rad) was used. Evaluation and quantification of the band intensities was done with the ImageLab software (Bio-Rad Laboratories).

### Superoxide dismutase activity

The SOD activity was measured with the direct spectrophotometric method using potassium superoxide as previously described, except that the assay was carried out at pH 9.5 (Marklund, 1976). The CSF samples were analyzed in duplicates and 20 µL of the original or antibody treated CSF samples were used for each measurement. For the measurement in blood samples, erythrocyte hemolysates were analyzed in duplicates and the activity related to the hemoglobin content (Andersen et al., 1998; Marklund, 1976).

### Total SOD1 ELISA

The total SOD1 protein content was determined by total SOD1 enzyme-linked immunosorbent assay (ELISA) essentially as previously described (Zetterstrom et al., 2011) but the secondary antibody was the chicken anti-SOD1 antibody described above, and as tertiary antibody a HRP-conjugated rabbit anti-Chicken IgY (Abcam, Cambridge, UK) was used. To assess the antibody preference for SOD1 folding state the chicken anti-SOD1 antibody was incubated with 5 µg/mL native and denatured SOD1. The native SOD1 was purchased from Sigma and denatured by a treatment with guanidinium chloride and EDTA as described (Zetterstrom et al., 2011). Two sequential incubations of the native SOD1 with antibody were carried out to ascertain that small amounts of non-native SOD1 present in the preparation did not influence the experiment. The two incubations resulted in equal amounts of SOD1 bound. The capacity of the chicken antibody to bind native SOD1 was 4-fold higher than its capacity to bind denatured SOD1 (Figure S1a). The primary rabbit anti-SOD1 antibody has, in a similar way, been found to have equal preference for native and denatured SOD1 (Zetterstrom et al., 2011).

### Statistics

Statistical analyses were done with the SPSS software (SPSS Inc., Chicago, Illinois, USA) version 29. All variables showed a normal distribution (Shapiro-Wilk test) except Erythrocyte SOD1 activity and CSF total protein. ANOVA with Bonferroni post hoc correction was used to search for differences between the groups and the skewness and kurtosis was acceptable for the Erythrocyte SOD1 activity to allow ANOVA. For the total protein and CSF SOD1 activity/total protein ratio, the non-parametric Kruskal-Wallis test with Bonferroni correction was used. Correlations were assessed by linear regression and Pearson’s correlation coefficient or Spearman’s rho. The significance level was set to 0.05. For a full statistical report see Table S2.

## Results

### A novel method for specific analysis of SOD1 activity in CSF

The cytosolic SOD1 and secretory SOD3 are evolutionary related Cu– and Zn-containing proteins with homologous active sites (Antonyuk et al., 2009) and there is no known inhibitor that can distinguish between them (Marklund, 1984). They are both, unlike the mitochondrial Mn-containing SOD2, efficiently inhibited by cyanide (Marklund et al., 1982). Cyanide-resistant SOD activity in human CSF is very low, however, around 0.2% of the total SOD activity (Jacobsson et al., 2001). To allow specific analysis of SOD1 activity in CSF, we here used immobilized anti-SOD3 antibodies to immunocapture the SOD3. The SOD activity was determined in CSF samples before and after an immunocapture step that depletes all the SOD3 from the sample. The SOD activity in the sample after immunocapture of SOD3 represents the SOD1 activity, and the difference between the first and the second measurements represents the SOD3 activity. As a proof-of-concept, we also designed a reversed assay with anti-SOD1-antibodies (Figure 1a).

**Figure 1.**
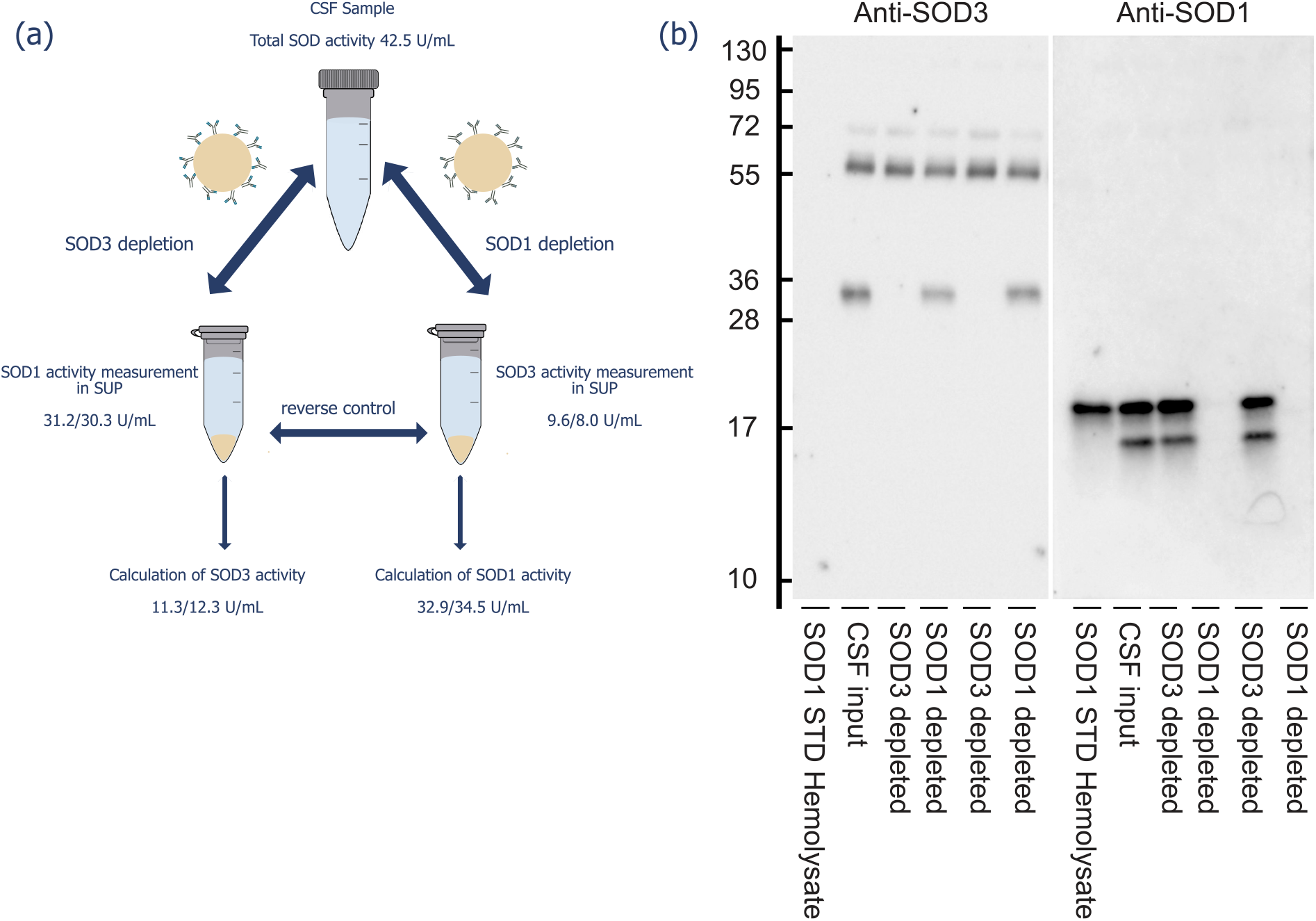
Specific determination of SOD1 activity in CSF. a) schematic illustration of immunocapture for determination of the enzymatic activity for the SOD1 and SOD3 isoforms in CSF. b) Western immunoblot showing the complete immunocapture of SOD3 or SOD1 in CSF that are fundamental for the presented method. Immunoblots stained for SOD3 (left) shows that there is no remaining SOD3 in CSF samples after depletion of SOD3 with immobilized anti-SOD3 antibodies and staining for SOD1 (right) shows no remaining SOD1 after depletion of SOD1 with immobilized anti-SOD1.

The anti-SOD3 or anti-SOD1 antibodies, in excess to the amount of SOD3 or SOD1 present in the CSF sample, were immobilized on CNBr-activated Sepharose and incubated. The Sepharose was then separated from the remaining SOD proteins in the soluble fraction by centrifugation (see method section for details). To determine the efficacy and optimal time for immunocapture, we evaluated different times from 1 hour up to 24 hours of binding using a pool of CSF collected from patients referred to the neurology ward at Umeå University Hospital for non-neurodegenerative causes. Already after two hours of capture with the anti-SOD3 antibody SOD3 was completely depleted from the sample, but all SOD1 remained in solution (Figure 2). Subsequently, two hours was chosen as binding time.

**Figure 2.**
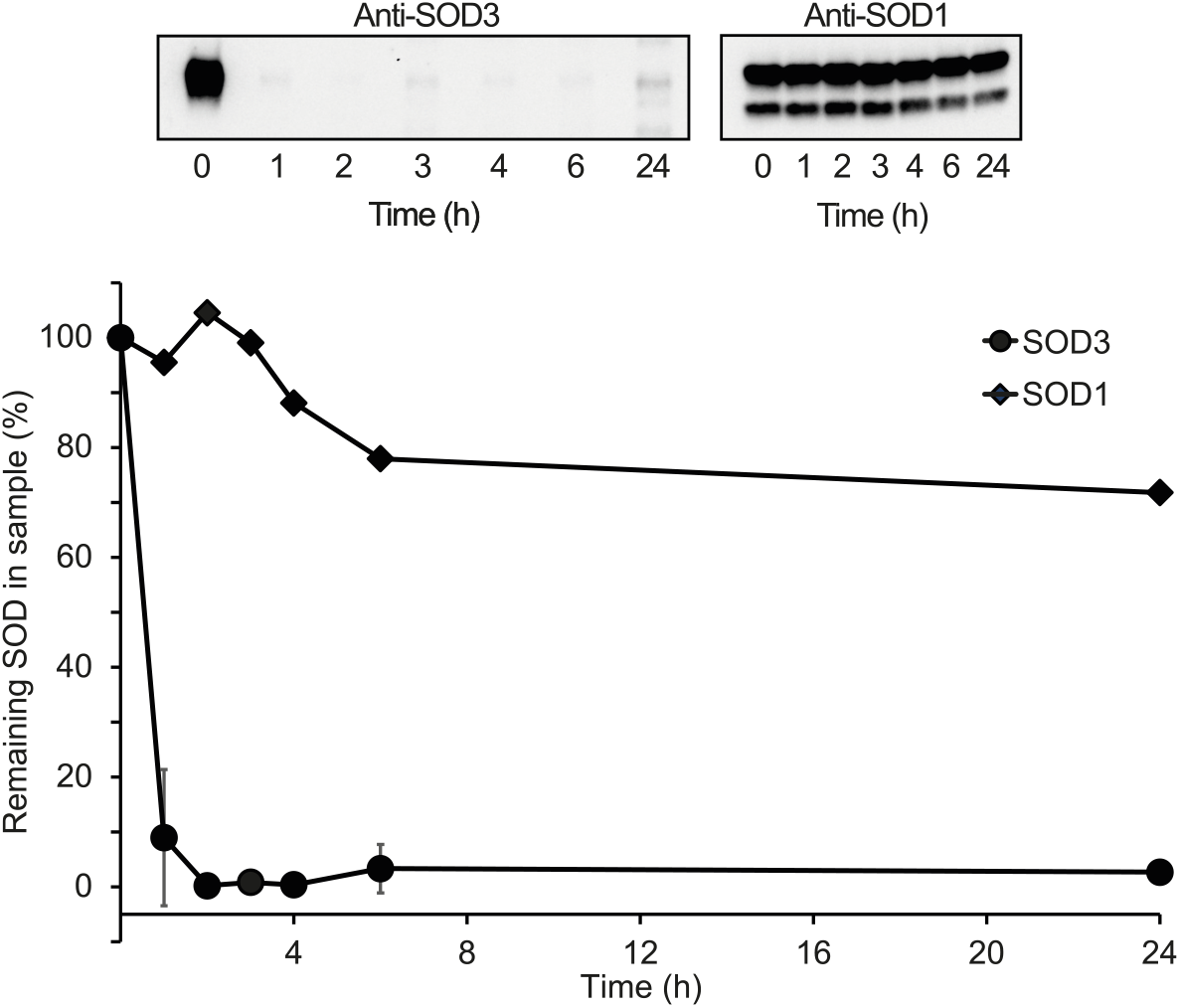
Time course for SOD3 immunocapture. SOD3 in a CSF sample was captured with immobilized anti-SOD3 antibodies for 1-24 hours. Remaining SOD3 and SOD1 in the sample was visualized by western immunoblot. Already after two hours virtually all SOD3 was captured but no effect on the content of SOD1 was seen. Top shows immunoblots and bottom quantification of the blots.

We next used the CSF pool for method evaluation. Aliquots of the CSF pool were assayed for SOD activity before and after immunocapture of SOD3 or SOD1. When anti-SOD3 antibodies were used, all SOD3 was depleted from the sample (shown by immunoblots for SOD3). The treatment did not affect the SOD1 content of the sample. The opposite pattern was apparent when the anti-SOD1 antibody was used (Figure 1b). Activity measurements before and after immunocapture rendered similar values for SOD1 activity independently of if anti-SOD3 or anti-SOD1 antibodies were used (Figure 1a). All further experiments were done using immunocapture of SOD3 for determination of SOD1 activity in CSF.

For precision determination, a similar pool of CSF from control individuals was aliquoted in n=25 separate samples and stored in –80°C. Aliquots were thawed, and the SOD activities determined in five aliquots on five consecutive days. The total SOD activity in the pool used was 36.2 ± 1.0 U/mL, the SOD1 activity was 26.5 ± 1.9 U/mL and the SOD3 activity was 9.7 ± 2.2 U/ml (mean ± standard deviation (SD)) giving relative SDs (cumulative variance (CV)%) of 2.9%, 7.2%, and 22.9% for total SOD activity, SOD1 activity, and SOD3 activity, respectively.

### SOD1 activity in CSF of ALS patients and controls

Using the novel method, we set out to measure SOD activities in CSF from controls, sporadic ALS patients without mutations in *SOD1* or *C9orf72*, ALS patients with mutations in *C9orf72* and ALS patients with mutations in *SOD1* (Table S1). The patients carrying *SOD1* mutations were stratified into stable and unstable mutants based on their SOD1 activities in erythrocytes (see material and methods).

All CSF samples were analyzed for total SOD activity, SOD1, and SOD3 activity, CSF SOD1 protein level, and total protein content. Erythrocyte SOD1 activity was measured at sample collection for the ALS patients and in 30 random control samples. These data are presented in Table 1 and Figure 3. Age at sampling and sample storage time for the different groups as well as the computed variables CSF SOD1/ erythrocyte SOD1, and CSF SOD1 specific activity are found in Table 1 and Figure S1.

**Figure 3.**
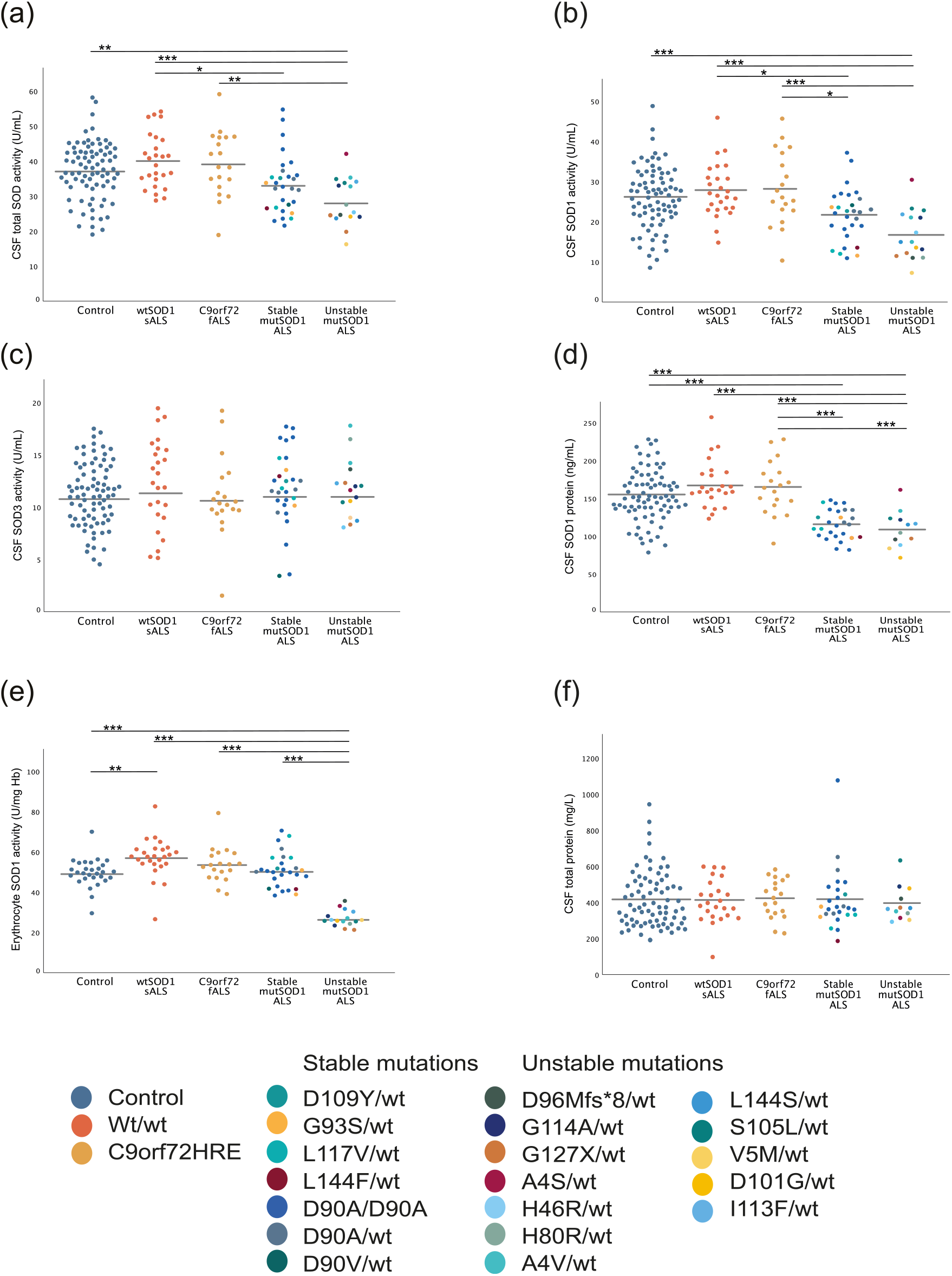
SOD activities and protein levels in CSF of ALS patients and controls. Scatter plots showing a) CSF total SOD activity, b) CSF SOD1 activity, c) CSF SOD3 activity, d) CSF SOD1 protein level, e) erythrocyte SOD1 activity, and f) CSF total protein. One dot represents one individual, bars represent means, for numerical values, se Table 1. Different colors are used for controls, sporadic ALS patients with wildtype SOD1, ALS patients with the *C9orf72* mutation, and different mutations in *SOD1*. Statistically significant differences are shown with * p<0.05; ** p<0.01, and *** p<0.001.

**Table 1.** Group characteristics and SOD activity and protein levels in CSF and erythrocytes of controls and ALS patients.

|  | Wildtype SOD1 |  |  | Mutant SOD1 |  | p |
| --- | --- | --- | --- | --- | --- | --- |
|  | Control<br>(n=81) | sALS<br>(n=26) | C9orf72 fALS<br>(n=19) | Stable SOD1 ALS<br>(n=29) | Unstable SOD1 ALS<br>(n=16) |  |
| Age at sampling (years) | 55.1 ± 15.6 | 60.7 ± 11.3 | 59.2 ± 11.2 | 55.6 ± 10.7 | 50.7 ± 13.7 | 0.127 |
| Storage time (years) | 9.1 ± 7.1 | 13.9 ± 6.1 | 9.7 ± 6.8 | 9.1 ± 9.1 | 6.0 ± 6.7 | 0.012 |
| CSF total SOD activity (U/mL) | 37.3 ± 8.2 | 40.1 ± 7.7 | 39.3 ± 9.3 | 33.3 ± 8.3 | 28.5 ± 6.8 | <0.001 |
| CSF SOD1 activity (U/mL) | 26.1 ± 7.6 | 27.8 ± 6.7 | 28.2 ± 9.1 | 21.6 ± 6.6 | 16.7 ± 6.1 | <0.001 |
| CSF SOD3 activity (U/mL) | 11.2 ± 3.0 | 12.3 ± 4.1 | 11.0 ± 3.8 | 11.8 ± 3.6 | 11.8 ± 2.8 | 0.594 |
| CSF SOD1 protein (ng/mL) | 119.3 ± 39.8 | 136.7 ± 36.5 (n=25) | 132.1 ± 36.5 | 74.9 ± 22.9 (n=27) | 66.9 ± 29.1 (n=12) | <0.001 |
| CSF total protein (mg/L) | 413 ± 155 (n=77) | 408 ± 119 (n=24) | 421 ± 107 | 415 ± 170 (n=26) | 393 ± 98 (n=12) | 0.844 |
| Erythrocyte SOD1 activity (U/mg Hb) | 49.6 ± 6.9 (n=29) | 57.1 ± 9.7 | 53.7 ± 8.8 | 51.1 ± 8.5 (n=28) | 27.0 ± 4.0 | <0.001 |
| CSF SOD1 specific activity (ng/U) | 4.5 ± 0.6 | 4.9 ± 0.5 (n=25) | 4.7 ± 0.6 | 3.6 ± 0.8 (n=27) | 4.0 ± 1.1 (n=12) | <0.001 |
| CSF SOD1 activity / Erc SOD1 activity | 0.54 ± 0.18 (n=30) | 0.51 ± 0.20 | 0.53 ± 0.17 | 0.43 ± 0.15 (n=27) | 0.62 ± 0.21 | 0.020 |
| CSF SOD1 activity / CSF total protein | 0.070 ± 0.032 (n=77) | 0.076 ± 0.034 (n=24) | 0.072 ± 0.029 | 0.055 ± 0.022 (n=26) | 0.045 ± 0.021 (n=12) | 0.002 |

Three control samples showed extremely high values (over 3 SD) for SOD activities in CSF as well as for total protein. None of these samples showed signs of gross hemolysis that could explain the high SOD activity. They were considered outliers and excluded from statistical evaluation. One control sample showed zero CSF SOD3 activity in repeated measurements. This sample was considered as an extreme outlier and excluded. Five control samples showed extremely high total protein content (over 3 SD) but normal CSF SOD activity. Only the total protein values were excluded for these five samples.

The CSF SOD1 activity levels increased with age (Figure S2b) in agreement with our previous studies (Jacobsson et al., 2001; Zetterstrom et al., 2011). The total protein levels likewise vary much and increase with age (Figure S2f) (Fautsch et al., 2023). The reason for the increase of total protein is not completely understood but may involve an age-dependent reduction in CSF turnover rates (Fautsch et al., 2023; Wichmann et al., 2021). Conceivably, the SOD1 activitiy variabilities might have partially the same background. We therefore calculated SOD1 activities divided by total protein in the CSF samples (Table 1). While the variabilities in these ratios were even larger than in SOD1 activities alone (Table 1), the increases with age disappeared (Figure 4a), suggesting that they are explained by reduced CSF turnover.

**Figure 4.**
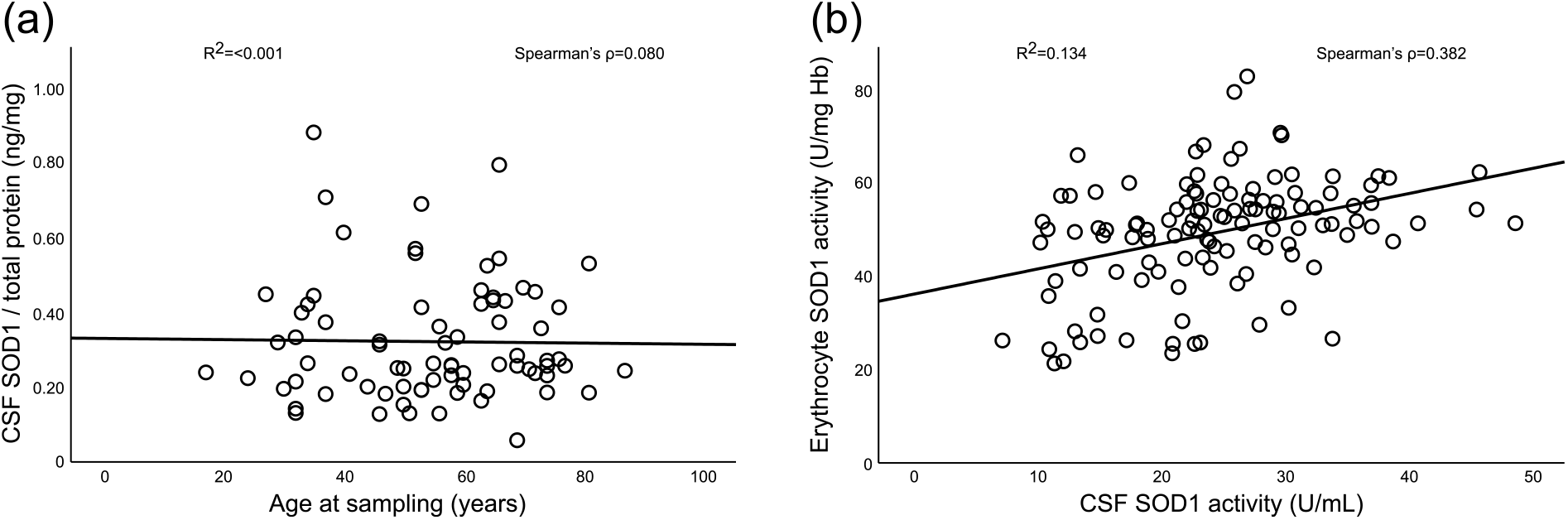
a) Correlation between the CSF SOD1 activity/CSF total protein ratio with age in the control individuals. No significant correlation was detected between this ratio and age (p=0.669). b) There is a significant correlation between the SOD1 activity in erythrocytes and CSF (p<0.001). Samples from both controls and ALS patients with and without SOD1 mutations are included to visualize the correlation in the full range of SOD1 activity found.

We also observed a significant increase of CSF SOD1 protein and CSF total SOD activity with increasing age (Figure S2e and a). However, no correlation for CSF SOD3 activity or erythrocyte SOD1 activity with increasing age was seen (Figure S2). Differences in age between the groups could therefore influence the CSF SOD activities, but since there was no significant difference in the age at sampling between the different groups (Table1, p=0.127) no corrections were needed.

### Large variation of SOD1 activity in CSF from ALS patients with *SOD1* mutations

The most common material for analysis of SOD1 activity in humans is erythrocytes due to the ease of sampling and the absence of other SOD isoenzymes than SOD1. Here we find minimally higher SOD1 activities in ALS patients without mutations in *SOD1* compared to the controls (Figure 3e), previously not commonly seen in blood (Bowling et al., 1995; Curti et al., 2002; Keskin et al., 2017). As expected, the ALS patients with unstable SOD1 variants showed significantly lower SOD1 activity in erythrocytes than all other groups (p<0.001, Figure 3e). In CSF the SOD1 activities show a different pattern. All three groups carrying wildtype *SOD1* have similar SOD1 activities that are significantly higher than the unstable mutant *SOD1* group (p<0.01, Figure 3b). The stable mutant SOD1 patients show significantly lower SOD1 activity than patients with wildtype *SOD1* (Figure 3b) but not compared to the controls (p=0.062). Normalization of the CSF SOD1 activity by the SOD1 activity in erythrocytes shows that the stable SOD1 mutants have lower than expected SOD1 activity in CSF compared to the unstable mutants (Figure S1c). The SOD1 activity in erythrocytes and CSF do correlate (Figure 4b, p<0.001). No significant differences regarding CSF SOD3 activity were found between any of the groups (Figure 3c). As described above, we found a large range in CSF SOD1 activity between different SOD1 mutants and similar differences have been found in blood (Keskin et al., 2017) and *in vitro* (Hayward et al., 2002).

Apart from the observed group level difference caused by *SOD1* mutations we also found a large variation among patients carrying the same *SOD1* mutation. For example, the patient with the highest SOD1 activity in the CSF was homozygous for the D90A mutation, but another homozygous D90A carrier showed the second lowest SOD1 activity of all patients analyzed with almost 4 times lower SOD1 activity in CSF. Several other mutants show a 2-fold difference in SOD1 activity in CSF from different individuals (Table S3). To further illustrate the large variability of the SOD1 activity in CSF, the control individuals showed a much higher variation in CSF with a CV% of 29% compared to 7% in erythrocytes (Table 1).

### Low ELISA reactivity of some mutant SOD1s

There was a discrepancy between results for SOD1 protein levels and activity in CSF for the mutant SOD1 group (Figure 3b and d). For the group with stable and unstable SOD1 mutations the ELISA-measured protein levels were 63% and 56% of the controls respectively. This is clearly lower than comparisons based on SOD1 activities which were 83% and 64% of the controls respectively (Table 1). ELISAs for SOD1 are typically based on two different antibodies raised against native wildtype SOD1. Mutant SOD1s, even if natively folded, may therefore show reduced reactivities in ELISA. In a previous study with ELISA we noted that the reactivity of native homozygous D90A SOD1 was 68% of the wildtype protein (Andersen et al., 1998) meaning that the lower protein levels observed in the SOD1 mutants could be due to different antibody reactivities.

In the current version of our total SOD1 ELISA, the secondary antibody raised in chickens showed a higher preference for native SOD1 than previously used antibodies (Figure S1a) and (Zetterstrom et al., 2011). With this ELISA the SOD1 protein level in CSF for the D90A homozygous patients was 60% which is lower than previously observed (Jacobsson et al., 2001). The SOD1 activity was 88% of the controls. Notably, the specific activity of pure isolated fully Cu– and Zn-charged D90A mutant SOD1 is the same as that of the wildtype enzyme (Marklund et al., 1997). This suggest that the ELISA response of the D90A protein is reduced to 68% also with our current ELISA configuration. For the other stable mutant SOD1s, all heterozygotes, the SOD1 activity was 75% and the SOD1 protein 65% compared to the controls. The wildtype SOD1 in these samples is expected to contribute about 50% of control levels, suggesting that the ELISA response of the mutants is reduced to similar extent as that of homozygous D90A. Even in the unstable mutants was the measured SOD1 activity higher than the protein compared with controls, 64% and 56%, respectively.

To further examine this, we measured the SOD1 protein content in erythrocytes with the SOD1 ELISA in a subset of the controls and patients with stable SOD1 mutations from whom samples were available. In erythrocytes, the D90A homozygous patients had 97% of the SOD1 activity (Figure 5a) and 66% of the SOD1 protein (Figure 5b), again showing 68% ELISA response. In the other stable SOD1 mutants, all heterozygotes, the SOD1 activity was 104% (Figure 5a) and the SOD1 protein 86% (Figure 5b). Accounting for the expected contribution of wildtype SOD1, this again suggests reductions in ELISA response similar to that of homozygous D90A.

**Figure 5.**
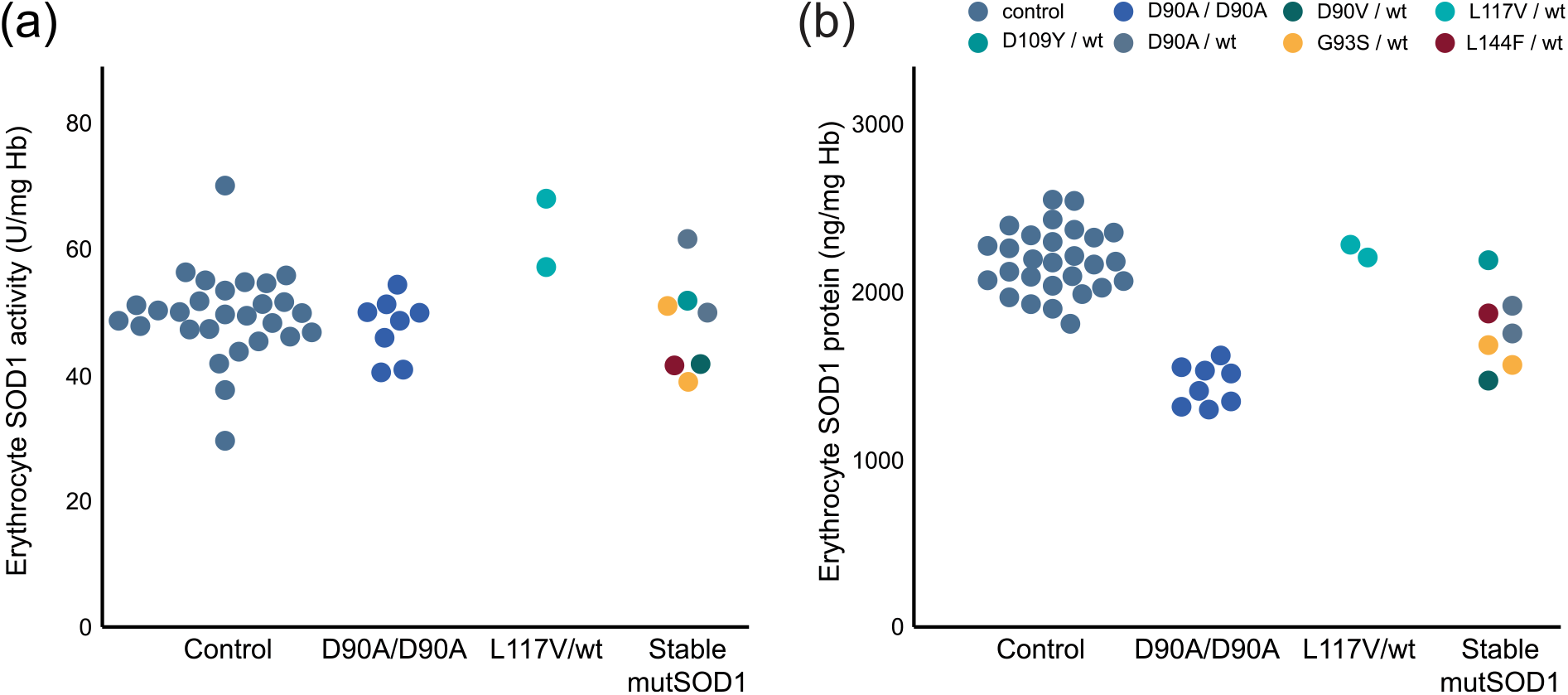
SOD1 activities and protein levels in erythrocytes. A subset of controls and ALS patients with stable SOD1 mutations were chosen for analysis of SOD1 protein in erythrocytes. a) SOD1 activity in erythrocytes for the selected individuals. b) SOD1 protein in erythrocytes. The D90A/D90A patients and L117V/ wildtype patients are plotted separately. Mutant SOD1s can give low ELISA reactivity in both CSF (Figure 3b and e) and erythrocytes compared to activity measurements.

Finally, we investigated the SOD1 protein level in CSF by immunoblotting samples from 27 controls and 21 patients with stable SOD1 mutations. Here we used an antipeptide antibody raised in rabbits against amino acids 24-39 in human SOD1. This antibody has a high preference for denatured SOD1 (Zetterstrom et al., 2011) and no patient carries a mutation in this region making the antibody suitable for western blot. Quantification showed that the patients had 75% of the control SOD1 protein, in good agreement with the CSF SOD1 activity which was 78% of the controls (not shown). We conclude that mutant SOD1s may commonly show reduced responses in ELISA. Assessment of SOD1 enzymatic activities based on such data will lead to underestimations.

## Discussion

Since it is not possible to collect biopsies of central nervous system (CNS) from ALS patients, the best estimate of CNS protein content in living humans is the CSF. We here present a new method to specifically quantify SOD1 enzymatic activity in CSF. This analysis is not trivial since the total SOD activity in CSF is low (Jacobsson et al., 2001; Marklund et al., 1982), constituting only around 0.2% of the SOD activity in CNS tissues and erythrocytes (Jonsson et al., 2009). Analysis of SOD activities in this range requires a sensitive method to get reliable results and the direct enzymatic method used here is around 30 times more sensitive than other commonly used methods (Beauchamp & Fridovich, 1971). By using a direct method for quantification of the SOD1 activity we can measure these low SOD activities that are found in the CSF with high precision.

The other complicating factor for SOD1 activity assessment in CSF is the presence of SOD3. While the active site cleft is structurally similar between SOD1 and SOD3 (Antonyuk et al., 2009), SOD3 is a much larger protein and they show significant sequence differences (Hjalmarsson et al., 1987). By using isoform-specific antibodies we could immunocapture all SOD3 in CSF samples and the remaining SOD activity accounts for the SOD1 activity. The procedure shows good precision, sensitivity, and reproducibility.

We measured the SOD1 activity in 171 CSF samples. In controls, we find that ca. 70% of the total SOD activity in CSF is SOD1 (Table 1) in accordance with previous estimates (Jacobsson et al., 2001; Strand & Marklund, 1992). The main superoxide defense scavenging system in the CSF is hence not the extracellular SOD3 detached from the heparan sulphate proteoglycans on cell surfaces but SOD1. The wildtype SOD1 protein in CSF was highly active with specific activities in the groups (Table 1) close to that of isolated native fully Cu– and Zn-charged SOD1 (≈4.3 ng/U) (Andersen et al., 1998; Marklund et al., 1997).

ALS patients with *SOD1* mutations show lower SOD1 activities in CSF compared to ALS patients with wildtype SOD1. Patients with unstable SOD1 variants show the lowest values (Table 1 and Figure 3b). There is no difference in SOD3 levels across all groups. Quantification of the SOD1 protein by ELISA shows a similar pattern as the activity measurements but the stable SOD1 variants show lower than expected protein levels (Table 1, Figure 3d). There is a larger discrepancy in the CSF than in erythrocytes (Figure 5). Erythrocytes have lifespans of around 120 days. Disordered proteins are efficiently recognized for degradation and there is no synthesis of new protein. As a result, the amounts of mutant SOD1 in erythrocytes vary widely depending on the degrees of destabilization, from close to the wildtype protein to <0.1% (Ezer et al., 2022; Jonsson et al., 2002; Park et al., 2023; Sato et al., 2005). The remaining mutant SOD1 protein is likely to mostly be active, resulting in SOD1 activities in heterozygotes varying from half to equal of controls (Keskin et al., 2017). Although weak, there is an overall correlation between the SOD1 activities in CSF and erythrocytes, but many patients deviate (Figure 4b). There are several explanations for the rather low concordance. First, the variability in CSF SOD1 activity is large among individuals, with CV% around 30% (Table 1). Second, the proteins are continuously replenished, and the turnover is more rapid in the CNS (≈25 days (Crisp et al., 2015)) expected to result in relatively higher levels of unstable variants. Third, the cellular matrix in CSF and blood is different with possible differences in recognition for protein degradation.

The mechanisms by which the normally intracellular SOD1 enters the CSF is not well understood. There is some evidence that SOD1 can be secreted (Cruz-Garcia et al., 2018; Hosomi et al., 2023; Mondola et al., 1996; Urushitani et al., 2006), but there may also be more unspecific leakage over the plasma membrane. Blood plasma contains only about 3 U/ml SOD1, meaning that leakage over the BBB should be an insignificant source (Marklund et al., 1986). We previously showed that in acute injury such as ischemic stroke there is increased cellular leakage leading, on average, to doubled SOD1 activities in CSF (Strand & Marklund, 1992). Here we find the SOD1 activities in CSF from ALS patients without *SOD1* mutations to be non-significantly higher than in controls, in accord with the much more slowly developing injury to cells in the motor system in ALS.

Even though reductions of SOD1 activity is not considered to be a primary cause of ALS, the SOD1 activity could still modify the course of ALS (reviewed in (Saccon et al., 2013)). Further emphasizing the importance of SOD1 activity and indicating vulnerability to low SOD1 levels in the CNS is the recent discovery of children homozygous for *SOD1* mutations that result in total-loss-of enzymatic function. These children are born normal but develop, from about the age of 6 months, the Infantile SOD Deficiency Syndrome (ISODDES) with progressive affection of both the upper and lower motor neuron systems but also with additional neurological and hematological abnormalities (Andersen et al., 2019; Çakar et al., 2023; de Souza et al., 2021; Dogan et al., 2024; Ezer et al., 2022; Park et al., 2023). So far, homozygosity for four *SOD1* mutations has been associated with causing ISODDES. The children with ISODDES suggest that there is a lower limit for SOD1 activity in the CNS below which serious neurological damage may develop (Andersen et al., 2019; Ezer et al., 2022; Park et al., 2023). We do not know where that limit is but somewhere below 50% of the normal level since there are *SOD1* mutation carriers with inactive forms (e.g. A4V, I113T) that live into the eight decades without showing symptoms of ISODDES or ALS (Cudkowicz et al., 1997). Patients with unstable SOD1 variants show around 50% SOD1 activity in erythrocytes but 60% in CSF (Table 1). Since current anti-SOD1 agents also affect the wildtype allele, the SOD1 activity could potentially be very low in the CNS of treated patients with inactive mutations. However, this study shows that also patients with apparently active forms of SOD1 in erythrocytes (Jonsson et al., 2002) and *in vitro* (Hayward et al., 2002) studies, do show decreased activity in CSF. Hence, all patients may be at risk of unhealthily low SOD1 activity in the CNS if chronically treated with total SOD1-lowering drugs.

This study also suggests that monitoring the degree of SOD1 reduction solely by assaying protein levels may be problematic. Methods based on anti-SOD1-antibodies can react differently to wildtype and mutant SOD1 and are difficult to standardize. Analysis of the SOD1 activity in CSF is a better alternative since it is the loss of enzymatic activity that likely drives ISODDES. Still, the variation in CSF SOD1 activity within the control individuals and ALS patients without *SOD1* mutations is very large, with CV% being 24-32% (Table 1). In comparison, the SOD1 activities in various parts of the CNS differ much less between individuals, with CV% around 13% in both controls and deceased D90A-homozygous ALS patients (Jonsson et al., 2009). The variabilities in erythrocytes of controls and ALS patients without SOD1 mutations also show CVs around 15%, as reported previously (Keskin et al., 2017). This so far unexplained variability in CSF may conceivably raise issues in longitudinal monitoring of the SOD1 activity in patients on SOD1-protein lowering therapies. A limitation of the present study is the lack of longitudinal CSF samples from individual patients for SOD1 activity measurements. However, it has been found that SOD1 protein levels in CSF measured with ELISA is rather constant in individuals and varies little over time (Lange et al., 2017; Winer et al., 2013).

If there is a lower limit for healthy SOD1 expression, this calls for treatment strategies to complement lowering *SOD1* expression. One would be to use mutant-specific ASO and siRNA’s but would prove difficult due to the large number of *SOD1* mutations, the 60+ mutations that are variants of unknown significance, the possibility of de novo mutations occurring (Müller et al., 2022), and urgency of early treatment, as well as economic considerations. Reduced expression of SOD1 slows down the prion formation by reducing the building blocks for the prion build up. Stopping the prion aggregation of SOD1 can also be done by removal of the SOD1 prions with e.g. anti-SOD1 antibodies (Lehmann et al., 2020; Maier et al., 2018; Urushitani et al., 2007). Such strategies may complement each other and likely more than one drug will be needed for efficient treatment of ALS.

## Author contributions

**Laura Leykam**: Investigation; visualization; writing – original draft; writing – review and editing **Karin M.E. Forsberg:** Resources; writing – review and editing **Ulrika Nordström**: Investigation; writing – review and editing **Karin Hjertkvist:** Investigation **Agneta Öberg:** Investigation **Eva Jonsson:** Investigation **Peter M. Andersen:** Conceptualization; resources; formal analysis; writing – review and editing; funding acquisition **Stefan L. Marklund:** Conceptualization; methodology; supervision; formal analysis; writing – review and editing **Per Zetterström:** Project administration; formal analysis; visualization; writing – original draft; writing – review and editing; funding acquisition

## Acknowledgments

We are indebted to the patients and their families for their help with this project. The authors thank Caitlin Henne for critically reading the final manuscript.

## Conflict of interest

LL, UN, KH, AÖ, EJ, SLM, and PZ: nothing to declare

KMEF: Clinical trial site investigator for Amylyx, Biogen, Ionis Pharmaceuticals, PTC-Pharmaceuticals, Sanofi and ITB-Med.

PMA: Paid consultancies and serve/have served on advisory boards for Biogen, Roche, Arrowhead, Avrion, Regeneron, uniQure, Voyager and Orphazyme A/S; clinical trial site investigator for AB Science, AL-S Pharma and Lilly, Amylyx, Alexion Pharmaceuticals, Biogen Idec, IBT-Med, IONIS Pharmaceuticals, Orion Pharma, PTH Pharmaceuticals, Sanofi. Since 1993 Director of the ALS-genetic laboratory at Umeå University Hospital that performs not-for-profit research genetic testing and free genetic testing including for SOD1. Since 2021, member of the ClinGen ALS Gene variant Curation Expert panel. External advisor to the European Medicine Agency

## Data availability statement

The data that support the findings of this study are available from the corresponding author upon reasonable request.

## Abbreviations

SOD1: superoxide dismutase-1
ALS: amyotrophic lateral sclerosis
ISODDES: Infantile SOD1 Deficiency Syndrome
CSF: cerebrospinal fluid
SOD3: extracellular superoxide dismutase
ASO: antisense oligonucleotides
EDTA: ethylenediaminetetraacetic acid
RT: room temperature
ELISA: enzyme-linked immunosorbent assay
SD: standard deviation
CV: cumulative variance
CNS: central nervous system

