## Supplementary information for "SOD1 Enzymatic Activity in CSF from ALS patients with and without *SOD1* mutations"

**SE-901 85 Umeå, SWEDEN**

**Table S1** Diagnosis of the Control samples

| Control Subject # | |
|---|---|
| 1 | Ehlers-Danlos Syndrome |
| 2 | Parestesia of unknown etiology |
| 3 | Cerebellar ataxia |
| 4 | Papilar edema of unknown etiology |
| 5 | Vertgo |
| 6 | Migraine attack |
| 7 | Polyneuropathy |
| 8 | Parestesia of unknown etiology |
| 9 | Parestesia of unknown etiology |
| 10 | Parestesia of unknown etiology |
| 11 | Seizure of unknown etiology |
| 12 | Lumbago |
| 13 | Glioblastoma |
| 14 | Sarcoidosis |
| 15 | Lumbar radiculitis of unknown etiology |
| 16 | Aphasia caused by cerebral hypoperfusion |
| 17 | Epilepsia |
| 18 | Headache |
| 19 | Sinuitis with headache |
| 20 | Polyneuropathy |
| 21 | Myalgia of unknown eitiology |
| 22 | headache |
| 23 | Acqvaductal stenosis with hydrocephalus |
| 24 | Hypesthesia of unknown etiology |
| 25 | Opticus neuritis |
| 26 | Intracraniel hypertension, obesitas |
| 27 | Myelopathy |
| 28 | Alzheimers Disease (MCI) |
| 29 | Myelopathy (post-ischemic?) |
| 30 | Healthy |
| 31 | Sarcoidosis |
| 32 | Migrain with aura |
| 33 | Cerebellar ataxia |
| 34 | Tension-type headache |
| 35 | Encefalitis |
| 36 | Headache |
| 37 | Headache |
| 38 | Encefalitis |
| 39 | Headache |
| 40 | Hydrocephalus with shunt infection |
| 41 | Headache |
| 42 | Headache |
| 43 | Muscle cramps of unknown etiology |
| 44 | Headache |

45 Sarcoidosis
46 Polyneuropathy
47 Headache
48 Polyneuropathy
49 Sarcoidosis?
50 Cerebral Vasculitis
51 AML with Acute Inflammatory Demyelinating Polyneuropathy
52 Myelitis
53 *Excluded from statistical analysis.* Amyloid angiopathy
54 Confusion (temporary)
55 Fasciculations of unknown etiology
56 Cancer of the prostatae
57 Headache
58 Migraine without aura
59 Mild neuropathy of unknown etiology
60 Sarcoidosis?
61 Myalgia of unknown etiology
62 *Excluded from statistical analysis.* Normal Pressure Hydrocephalus
63 Parkinsonism
64 Neuropathy
65 Polyneuropoathy (drug induced)
66 Parkinsons Disease
67 Cerebellar Ataxia
68 Spastic hemiparesis
69 Acute Myeloid Leukemia
70 Encephalomyelitis (autoimmune)
71 Headache
72 Seizure of unknown etiology
73 Reumatism
74 *Excluded from statistical analysis.* Progressive suprenuclear paresis
75 headache
76 psychosis
77 Chronic walking difficulties of unknown etiology
78 Parkinson's Disease
79 III Crannial nerve paresis of unknown etiology
80 Cervical myelitis
81 Normal pressure hydrocephalus
82 Nystagmus
83 Polyneuropathy
84 *Excluded from statistical analysis.* Stroke and miild polyneuropathy
85 Cerebral vasculopathy

Final diagnosis of the 85 control subjects. 41 were women.
Median age at sampling 57 ± 15 years.
Median storage time: 4.7 years.

**Table S2** Full statistical report

**ANOVA**

| | | Sum of Squares | df | Mean Square | F | Sig. |
|---|---|---|---|---|---|---|
| Age at sampling (years) | Between Groups | 1290.642 | 4 | 322.660 | 1.820 | 0.127 |
| | Within Groups | 29422.516 | 166 | 177.244 | | |
| | Total | 30713.158 | 170 | | | |
| Storage time (years) | Between Groups | 705.100 | 4 | 176.275 | 3.315 | 0.012 |
| | Within Groups | 8825.962 | 166 | 53.168 | | |
| | Total | 9531.062 | 170 | | | |
| CSF Total SOD activity (U/mL) | Between Groups | 1847.738 | 4 | 461.934 | 6.954 | <0.001 |
| | Within Groups | 11027.027 | 166 | 66.428 | | |
| | Total | 12874.765 | 170 | | | |
| CSF SOD1 activity (U/mL) | Between Groups | 1923.893 | 4 | 480.973 | 8.823 | <0.001 |
| | Within Groups | 9049.353 | 166 | 54.514 | | |
| | Total | 10973.246 | 170 | | | |
| CSF SOD3 activity (U/mL) | Between Groups | 31.791 | 4 | 7.948 | 0.698 | 0.594 |
| | Within Groups | 1890.386 | 166 | 11.388 | | |
| | Total | 1922.177 | 170 | | | |
| CSF SOD1 protein (ng/mL) | Between Groups | 88696.981 | 4 | 22174.245 | 16.531 | <0.001 |
| | Within Groups | 213275.412 | 159 | 1341.355 | | |
| | Total | 301972.393 | 163 | | | |
| Erc SOD1 activity (U/mg Hb) | Between Groups | 10027.921 | 4 | 2506.980 | 38.847 | <0.001 |
| | Within Groups | 7292.387 | 113 | 64.534 | | |
| | Total | 17320.308 | 117 | | | |
| CSF SOD1 specific activity (ng/U) | Between Groups | 29.948 | 4 | 7.487 | 17.759 | <0.001 |
| | Within Groups | 67.032 | 159 | 0.422 | | |
| | Total | 96.980 | 163 | | | |
| CSF SOD1 activity / Erc SOD1 activity | Between Groups | 0.397 | 4 | 0.099 | 3.035 | 0.020 |
| | Within Groups | 3.667 | 112 | 0.033 | | |
| | Total | 4.065 | 116 | | | |

**Pairwise Comparisons**

**CSF total protein (mg/L)**

| Sample 1-Sample 2 | Test Statistic | Std. Error | Std. Test Statistic | Sig. | Adj. Sig.[a] |
|---|---|---|---|---|---|
| Unstable mutSOD1 ALS-control | 1.739 | 14.200 | 0.122 | 0.903 | 1.000 |
| Unstable mutSOD1 ALS-Stable mutSOD1 ALS | 2.737 | 15.968 | 0.171 | 0.864 | 1.000 |
| Unstable mutSOD1 ALS-wtSOD1 sALS | 8.604 | 16.177 | 0.532 | 0.595 | 1.000 |
| Unstable mutSOD1 ALS-C9orf72 fALS | 12.294 | 16.871 | 0.729 | 0.466 | 1.000 |
| control-Stable mutSOD1 ALS | -0.998 | 10.378 | -0.096 | 0.923 | 1.000 |
| control-wtSOD1 sALS | -6.865 | 10.697 | -0.642 | 0.521 | 1.000 |
| control-C9orf72 fALS | -10.555 | 11.721 | -0.901 | 0.368 | 1.000 |
| Stable mutSOD1 ALS-wtSOD1 sALS | 5.867 | 12.952 | 0.453 | 0.651 | 1.000 |
| Stable mutSOD1 ALS-C9orf72 fALS | 9.557 | 13.810 | 0.692 | 0.489 | 1.000 |
| wtSOD1 sALS-C9orf72 fALS | -3.690 | 14.050 | -0.263 | 0.793 | 1.000 |

Each row tests the null hypothesis that the Sample 1 and Sample 2 distributions are the same.

Asymptotic significances (2-sided tests) are displayed. The significance level is 0.050.

a. Significance values have been adjusted by the Bonferroni correction for multiple tests.

**Pairwise Comparisons**

**CSF SOD1 activity / CSF total protein**

| Sample 1-Sample 2 | Test Statistic | Std. Error | Std. Test Statistic | Sig. | Adj. Sig.[a] |
|---|---|---|---|---|---|
| Unstable mutSOD1 ALS-Stable mutSOD1 ALS | 22.026 | 15.968 | 1.379 | 0.168 | 1.000 |
| Unstable mutSOD1 ALS-control | 43.139 | 14.200 | 3.038 | 0.002 | 0.024 |
| Unstable mutSOD1 ALS-C9orf72 fALS | 51.175 | 16.871 | 3.033 | 0.002 | 0.024 |
| Unstable mutSOD1 ALS-wtSOD1 sALS | 52.875 | 16.177 | 3.269 | 0.001 | 0.011 |
| Stable mutSOD1 ALS-control | 21.113 | 10.378 | 2.034 | 0.042 | 0.419 |
| Stable mutSOD1 ALS-C9orf72 fALS | 29.150 | 13.810 | 2.111 | 0.035 | 0.348 |
| Stable mutSOD1 ALS-wtSOD1 sALS | 30.849 | 12.952 | 2.382 | 0.017 | 0.172 |
| control-C9orf72 fALS | -8.037 | 11.721 | -0.686 | 0.493 | 1.000 |
| control-wtSOD1 sALS | -9.736 | 10.697 | -0.910 | 0.363 | 1.000 |
| C9orf72 fALS-wtSOD1 sALS | 1.700 | 14.050 | 0.121 | 0.904 | 1.000 |

Each row tests the null hypothesis that the Sample 1 and Sample 2 distributions are the same.

Asymptotic significances (2-sided tests) are displayed. The significance level is 0.050.

a. Significance values have been adjusted by the Bonferroni correction for multiple tests.

**Table S3** CSF SOD1 activities and erythrocyte SOD1 activities in ALS patients with mutant SOD1.

| SOD1 mutation | CSF SOD1 activity (U/ml) | n | Mean | SD | Rank CSF SOD1 | Erc SOD1 activity (U/mg Hb) | n | Mean | SD | Rank Erc SOD1 |
|---|---|---|---|---|---|---|---|---|---|---|
| **D90A/D90A** | 37.2 | 16 | 23.1 | 7.4 | 1 | 50.5 | 16 | 49.9 | 8.5 | 14 |
| | 35.2 | | | | 2 | 48.7 | | | | 18 |
| | 29.8 | | | | 4 | 70.6 | | | | 1 |
| | 27.3 | | | | 5 | 54.3 | | | | 8 |
| | 27,0 | | | | 6 | 40.4 | | | | 25 |
| | 26.7 | | | | 7 | 51.2 | | | | 11 |
| | 26.3 | | | | 8 | 38.3 | | | | 27 |
| | 22.1 | | | | 19 | 47.9 | | | | - |
| | 21.2 | | | | 21 | 48.6 | | | | 19 |
| | 19.1 | | | | 25 | 42.9 | | | | 21 |
| | 19,0 | | | | 26 | 47.9 | | | | 20 |
| | 19,0 | | | | 27 | 49.8 | | | | 17 |
| | 18.2 | | | | 28 | 51.2 | | | | 12 |
| | 16.5 | | | | 30 | 40.8 | | | | 24 |
| | 13.4 | | | | 35 | 65.8 | | | | 3 |
| | 10.9 | | | | 44 | 49.9 | | | | 16 |
| **A4S/wt** | 30.4 | 1 | | | 3 | 33.1 | 1 | | | 29 |
| **D90A/wt** | 25.7 | 5 | 22.9 | 1.8 | 9 | 57.5 | 4 | 55.3 | 5.3 | 5 |
| | 23.1 | | | | 14 | 61.6 | | | | 4 |
| | 22.7 | | | | 16 | - | | | | - |
| | 22.4 | | | | 18 | 50.1 | | | | 15 |
| | 20.8 | | | | 24 | 51.9 | | | | 9 |
| **D90V/wt** | 24.1 | 1 | | | 10 | 41.7 | 1 | | | 22 |
| **G93S/wt** | 23.6 | 2 | 17.6 | | 11 | 50.9 | 2 | 44.9 | | 13 |
| | 11.6 | | | | 40 | 38.9 | | | | 26 |
| **L117V/wt** | 23.6 | 3 | 16.1 | 6.5 | 12 | 67.9 | 3 | 60.7 | 6.2 | 2 |
| | 12.7 | | | | 37 | 57.1 | | | | 6 |
| | 12,0 | | | | 39 | 57.1 | | | | 7 |
| **S105L/wt** | 23.3 | 2 | 23.1 | | 13 | 25.7 | 2 | 25.6 | | 37 |
| | 22.8 | | | | 15 | 25.4 | | | | 39 |
| **D109Y/wt** | 22.7 | 1 | | | 17 | 51.8 | 1 | | | 10 |
| **I113F/wt** | 21.8 | 1 | | | 20 | 30.3 | 1 | | | 31 |
| **A4V/wt** | 21.1 | 2 | 18.1 | | 22 | 25.5 | 2 | 26.3 | | 38 |
| | 15,0 | | | | 31 | 27.1 | | | | 33 |
| **G114A/wt** | 21,0 | 2 | 17.1 | | 23 | 23.4 | 2 | 25.8 | | 41 |
| | 13.1 | | | | 36 | 28.1 | | | | 32 |
| **H46R/wt** | 17.3 | 1 | | | 29 | 26.2 | 1 | | | 34 |
| **L144S/wt** | 15,0 | 1 | | | 32 | 31.7 | 1 | | | 30 |
| **D101G/wt** | 13.6 | 1 | | | 33 | 25.8 | 1 | | | 36 |
| **L144F/wt** | 13.5 | 1 | | | 34 | 41.5 | 1 | | | 23 |
| **G127X/wt** | 12.2 | 2 | 11.9 | | 38 | 21.7 | 2 | 21.5 | | 42 |
| | 11.5 | | | | 41 | 21.3 | | | | 43 |
| **H80R/wt** | 11.1 | 1 | | | 42 | 24.3 | 1 | | | 40 |
| **D96Mfs*8/wt** | 11,0 | 1 | | | 43 | 35.7 | 1 | | | 28 |
| **V5M/wt** | 7.3 | 1 | | | 45 | 26.2 | 1 | | | 35 |

**Figure S1**

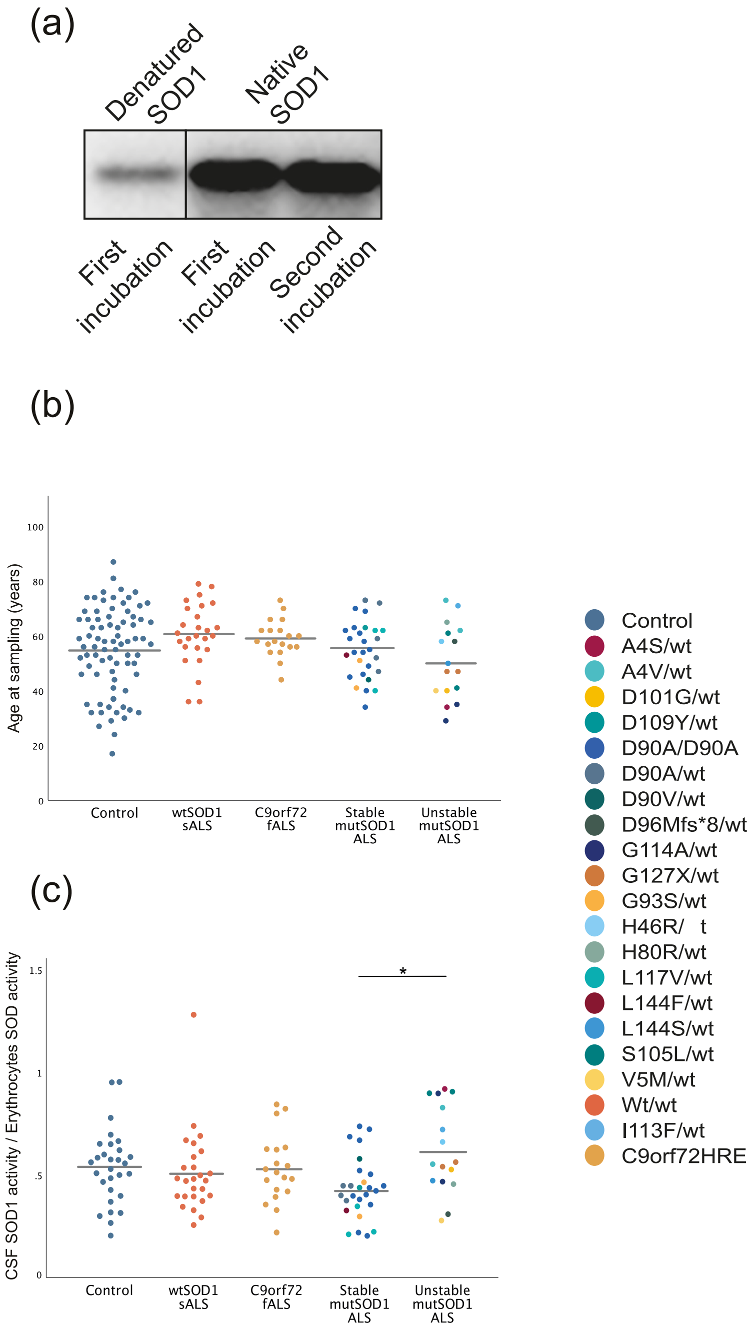
(a)
Denatured SOD1
Native SOD1
First incubation
First incubation
Second incubation
(b)
Age at sampling (years)
100
80
60
40
20
0
Control
wtSOD1 sALS
C9orf72 fALS
Stable mutSOD1 ALS
Unstable mutSOD1 ALS
(c)
CSF SOD1 activity / Erythrocytes SOD activity
1.5
1
.5
0
*
Control
wtSOD1 sALS
C9orf72 fALS
Stable mutSOD1 ALS
Unstable mutSOD1 ALS
Control
A4S/wt
A4V/wt
D101G/wt
D109Y/wt
D90A/D90A
D90A/wt
D90V/wt
D96Mfs*8/wt
G114A/wt
G127X/wt
G93S/wt
H46R/ t
H80R/wt
L117V/wt
L144F/wt
L144S/wt
S105L/wt
V5M/wt
Wt/wt
I113F/wt
C9orf72HRE

**Figure S2**

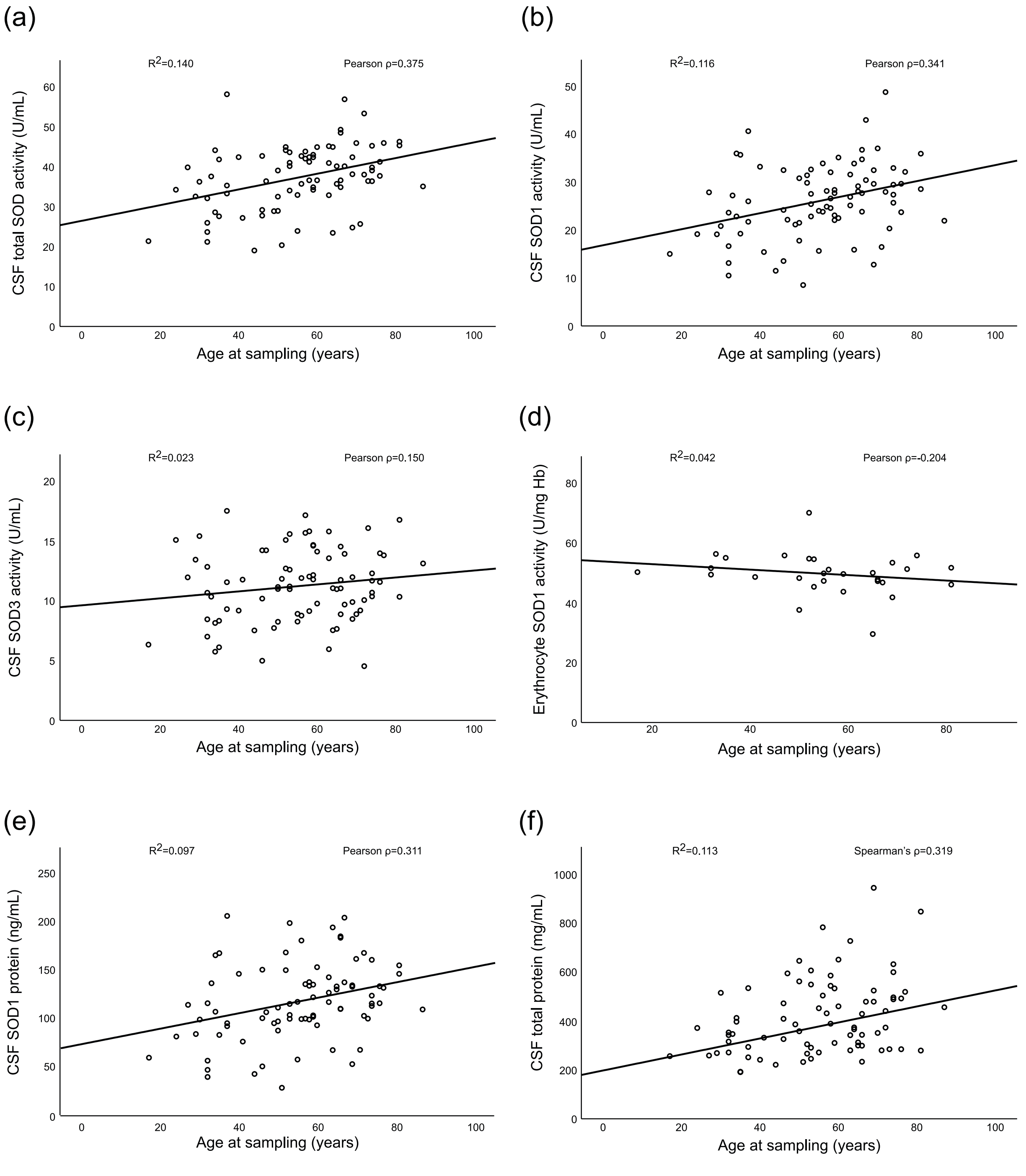
(a)
R2=0.140
Pearson ρ=0.375
CSF total SOD activity (U/mL)
Age at sampling (years)
(b)
R2=0.116
Pearson ρ=0.341
CSF SOD1 activity (U/mL)
Age at sampling (years)
(c)
R2=0.023
Pearson ρ=0.150
CSF SOD3 activity (U/mL)
Age at sampling (years)
(d)
R2=0.042
Pearson ρ=-0.204
Erythrocyte SOD1 activity (U/mg Hb)
Age at sampling (years)
(e)
R2=0.097
Pearson ρ=0.311
CSF SOD1 protein (ng/mL)
Age at sampling (years)
(f)
R2=0.113
Spearman's ρ=0.319
CSF total protein (mg/mL)
Age at sampling (years)

### Figure S3

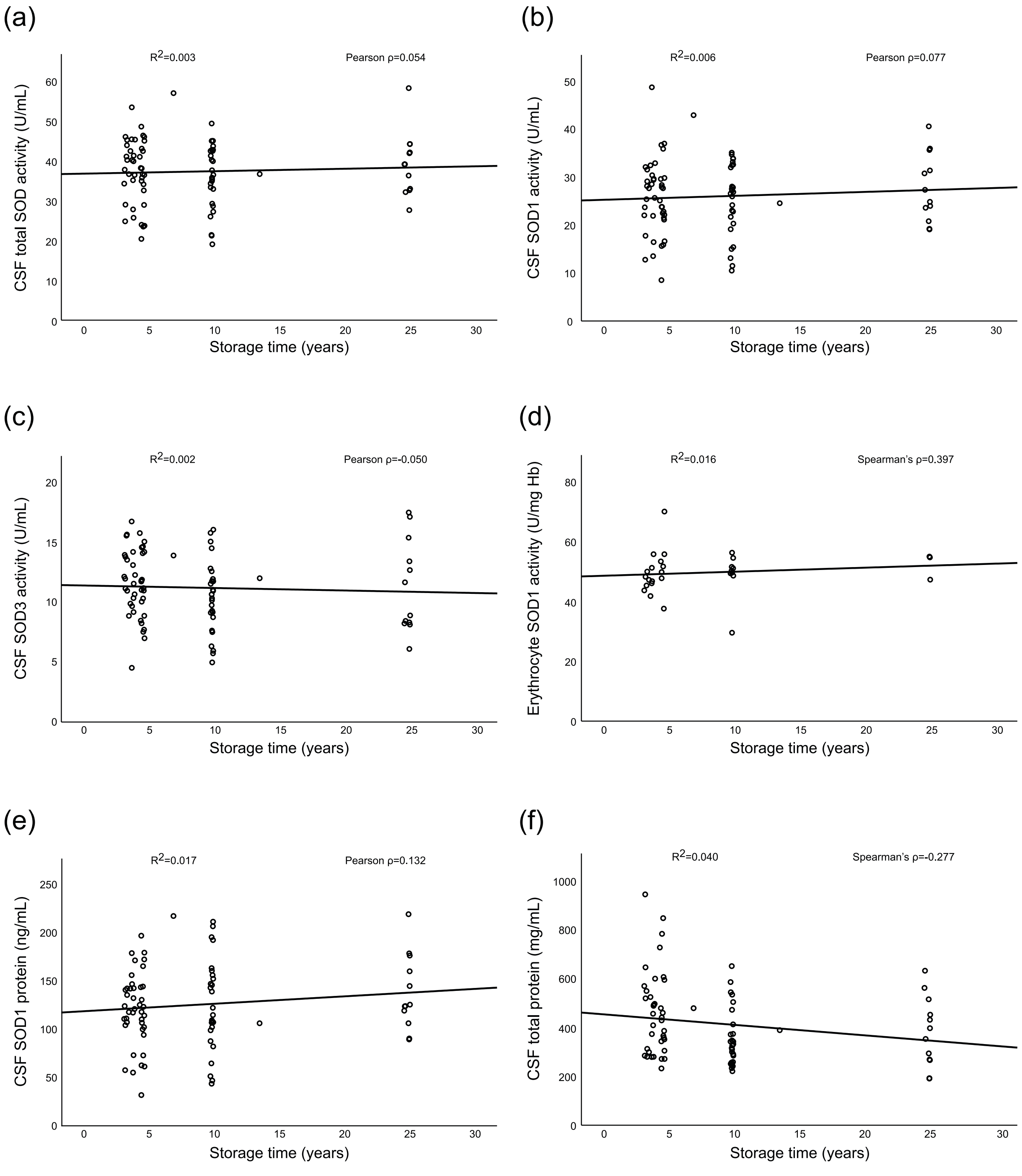
(a)
$R^2$=0.003
Pearson ρ=0.054
CSF total SOD activity (U/mL)
Storage time (years)
(b)
$R^2$=0.006
Pearson ρ=0.077
CSF SOD1 activity (U/mL)
Storage time (years)
(c)
$R^2$=0.002
Pearson ρ=-0.050
CSF SOD3 activity (U/mL)
Storage time (years)
(d)
$R^2$=0.016
Spearman's ρ=0.397
Erythrocyte SOD1 activity (U/mg Hb)
Storage time (years)
(e)
$R^2$=0.017
Pearson ρ=0.132
CSF SOD1 protein (ng/mL)
Storage time (years)
(f)
$R^2$=0.040
Spearman's ρ=-0.277
CSF total protein (mg/mL)
Storage time (years)

**Legends to supplementary figures:**

Figure S1 a) Immunocapture of native and denatured SOD1 with the chicken anti-SOD1 antibody used as secondary antibody in the SOD1 ELISA (see materials and methods). Scatter plots showing b) age at sampling and c) CSF SOD1 activity/erythrocyte SOD1 for ALS patients and controls. One dot represents one individual, bars represent means, for numerical values, see Table 1. Different colors are used for controls, sporadic ALS patients with wildtype SOD1, familial ALS patients with *C9orf72* mutations, and different mutations in *SOD1*. Statistically significant differences are shown with * $p<0.05$.

Figure S2 Effect of age at sampling on SOD activities in control individuals. a) CSF total SOD activity, b) CSF SOD1 activity, e) CSF SOD1 protein level, and f) CSF total protein are all positively correlated with age at sampling ($p<0.01$). CSF SOD3 activity (c) shows a weak positive correlation with increasing age and d) erythrocyte SOD1 activity shows a weak negative correlation with increasing age, however both unsignificant ($p>0.05$).

Figure S3 Effect of sample storage time on SOD activities in control individuals. f) CSF total protein show a negative correlation with sample storage time ($p=0.015$) and d) erythrocyte SOD1 activity show a weak positive correlation with sample storage time ($p=0.033$). a) CSF total SOD activity, b) CSF SOD1 activity, c) CSF SOD3 activity, and e) CSF SOD1 protein level show no significant correlation with sample storage time ($p>0.05$).
